# Adolescent Stress Exposure: Behavioral Consequences and Molecular Mechanisms in Corticolimbic Networks

**DOI:** 10.64898/2026.05.08.723933

**Authors:** E.M. Cotella, R.D. Moloney, P. Mahbod, S.E. Martelle, R.L. Morano, B.A. Packard, J.P. Herman

## Abstract

**Introduction:** Adolescence is a sensitive developmental period during which chronic stress can induce lasting adaptations in corticolimbic circuits involved in stress regulation, cognition, and emotional behavior. We examined the long-term behavioral, endocrine, and molecular consequences of adolescent chronic variable stress (CVS) in male and female rats, focusing on the infralimbic cortex (IL) and basolateral amygdala (BLA)

**Methods:** Sprague Dawley rats of both sexes were exposed to CVS during late adolescence and evaluated in adulthood after an extensive recovery period. Behavioral testing included cued fear conditioning and extinction recall, delayed spatial win-shift, novel object recognition, Morris water maze, three-chamber social behavior, and passive avoidance. HPA-axis reactivity to acute restraint was assessed. Targeted qPCR was used to measure stress-related gene expression in the IL and BLA immediately after stress or after a 5-week recovery period

**Results:** Adolescent CVS did not cause generalized cognitive impairment, but instead produced selective, sex-specific effects. Females had reduced HPA responses to acute stress and mild deficits in delayed spatial win-shift performance, together with long-term IL changes in genes related to adrenergic signaling, plasticity, and GABA clearance. Males showed enhanced Morris water maze probe retention, weaker novel object discrimination, altered passive avoidance with marked inter-individual variability, and enhanced social preference. At the molecular level, males exhibited long-term upregulation of Fkbp5 in IL and downregulation of PACAP, α1D adrenergic receptor, and proenkephalin in BLA, whereas females showed delayed PACAP upregulation in BLA

**Discussion:** Adolescent CVS induces persistent, sex- and region-specific recalibration of corticolimbic function, supporting distinct patterns of vulnerability and resilience, rather than uniform stress pathology.

## 2. Introduction

The study of factors influencing the adolescent brain holds significant translational relevance, particularly considering the elevated occurrence of first-episode onset affective conditions, such as depression and anxiety disorders, during this developmental phase (Solmi et al., 2022; Keyes and Platt, 2023a). Adolescence represents a critical period of substantial brain development (Larsen and Luna, 2018; Keyes and Platt, 2023b), with various brain regions undergoing maturation through processes of pruning, myelination, and dynamic shifts in the expression levels of multiple genes (Gogtay et al., 2004; Kwon et al., 2020; Dipnall et al., 2023; Fleming and McDermott, 2024; Peña, 2025). This developmental progression is integral to the establishment of proper adult physiological and behavioral responses, and disruptions to this process may have enduring effects on functions regulated by these brain regions.

Chronic environmental stressors can represent major disruptors of adolescent neurodevelopment. Previous studies show that stress during adolescence elicits distinct endocrine and behavioral responses (Romeo, 2010; Foilb et al., 2011; Jankord et al., 2011). Moreover, the long-term consequences of adolescent stress are not only age-specific (Cotella et al., 2019) but also sex-dependent (Smith et al., 2018; Cotella et al., 2020). Contrary to the widespread assumption that developmental stress exclusively produces negative outcomes, recent evidence, including work from our group, demonstrates that certain forms of adolescent stress can also promote resilience to later stressors (Kendig et al., 2011; Suo et al., 2013; Deng et al., 2017; Cotella et al., 2023).

Chronic stress robustly affects the structure and function of corticolimbic regions involved in behavioral and endocrine regulation, including the medial prefrontal cortex (mPFC) and amygdala (Amy) (Radley et al., 2015; Herman et al., 2020; Wellman et al., 2020). These regions work in concert with other interconnected networks to support cognitive processes such as goal-directed behavior, defensive responses, and learning and memory, all of which can be altered by stress exposure (McEwen, 2017). Indeed, the stress sensitivity of corticolimbic circuitry has been mechanistically linked to vulnerability for psychiatric disorders (Andersen and Teicher, 2008).

We recently identified mechanisms within the infralimbic subdivision of the mPFC that contribute to resilience in a PTSD-like model following adolescent stress (Cotella et al., 2023) in line with evidence that stress during adolescence can lead to distinct patterns of vulnerability and resilience. Building on this work, in the present study, we framed adolescent CVS as a developmental perturbation that may reorganize brain circuits and generate selective, rather than universally maladaptive, adult outcomes. We employed a series of behavioral tasks to characterize long-term consequences of our adolescent chronic variable stress (CVS) protocol, with particular emphasis on cognitive domains dependent on prefrontal and broader corticolimbic function. In parallel, we examined potential molecular substrates underlying these enduring effects by assessing candidate genes involved in stress regulation. This analysis centered on the infralimbic (IL) and basolateral amygdala (BLA), both key regions in top-down control of stress responses, fear expression, and defensive behavior (Gilmartin et al., 2014; Bloodgood et al., 2018; Marek et al., 2019).

## 3. Materials and Methods

### 3.1 Animals

Male and freely cycling female Sprague Dawley rats were bred in-house, weaned at postnatal day 21 (PND21) and pair-housed in standard clear cages (20 cm height x 22 cm width x 43 cm length) under a 12 h light/12 h dark cycle (lights on at 7:00 am). The room was maintained at a constant temperature of 23 ± 2 °C, with ad libitum access to food and water (unless specified by the experiment). All tests were conducted during the light cycle between 09:00 AM and 2:00 PM, except for the delayed spatial win shift test, for which the light cycle was reversed to test animals during their active cycle. All procedures and animal care were approved by the University of Cincinnati Institutional Animal Care and Use Committee.

### 3.2 Adolescent chronic variable stress (adol CVS)

Following weaning, animals were randomly assigned to experimental groups, with no more than two littermates in each group. Rats underwent our CVS protocol during late adolescence (Jankord et al., 2011). In Experiments 1 and 2, a 3-week CVS protocol commenced at PND 40 ± 2. In subsequent experiments, based on prior results indicating efficacy and sufficiency (Wulsin et al., 2016; Cotella et al., 2020), we adopted a 2-week CVS protocol starting at PND 46 ± 2 to prevent prolonged weight loss and mitigate potential metabolic combinatory effects of stress during pubertal development.

The CVS paradigm involved twice-daily application of unpredictable variable stressors (AM and PM): 1) 1 h shaker stress (100 rpm), 2) 1 h cold room (4 °C), 3) 30 min restraint in a plexiglass adjustable cylinder (not in Experiments 1 and 2), 4) 30 min hypoxia exposure (8 % O2 and 92 % N2), 5) 10 min group cold swim (17 ± 1 °C), 6) 20 min group warm swim (32 ± 1 °C). Additionally, animals experienced overnight stressors every two days: 1) Individual housing, 2) Social crowding (six rats per cage). Control animals were maintained in the same room and only handled for normal husbandry. All CVS-related procedures, except for overnight stressors, were conducted in a separate room. Following CVS, animals were allowed several weeks (see each experiment details) to recover into adulthood before undergoing the corresponding evaluations.

### 3.3 Methods

#### 3.3.1 Experiment 1

##### Fear Conditioning

Five weeks after the end of a 3-week adol CVS protocol, a group of rats was evaluated through an auditory cued fear conditioning paradigm. On day 1, rats were habituated to the chamber for 5 minutes and then received five 30-second tones (CS), each paired with a 0.5-second, 0.5 mA shock (US), with 3-minute intertrial intervals. On day 2, extinction was tested by re-exposing rats to the tone in the same chamber for 5 minutes, followed by 20 unreinforced CS presentations (ITI = 3 minutes). On day 3, extinction recall was assessed after 5 minutes of exploration, followed by three CS presentations (ITI = 3 minutes).

##### ACTH and Corticosterone assessment

The HPA response to acute restraint stress was assessed two weeks following fear conditioning. Rats were restrained for 30 minutes, and blood samples were collected from the tail at five time points: immediately before restraint (0 min), and at 15 and 30 minutes during restraint. Additional samples were collected at 60 and 120 minutes after the onset of restraint, corresponding to 30 and 90 minutes after the animals were released from the restraining tubes.

Blood collection was performed within 3 minutes by gently removing the clot from the distal tail after the initial clip. To minimize circadian hormonal variability, all procedures occurred during the circadian low of the diurnal corticosterone (CORT) cycle, with blood samples collected before 1:00 PM. After collection, blood was centrifuged at 3500 x g for 15 minutes at 4 °C, and plasma samples were stored at −20 °C until hormone levels were determined.

CORT and adrenocorticotropic hormone (ACTH) plasma levels were measured via radioimmunoassay, as previously described (Cotella et al., 2020). CORT concentration was determined using 125I RIA kits (MP Biomedicals Inc., Orangeburg NY), and ACTH was determined by radioimmunoassay using 125I ACTH as a trace (Amersham Biosciences) with ACTH antiserum (kindly donated by Dr. William Engeland, University of Minnesota) at a 1:120,000 dilution (Ostrander et al., 2006). The ratio of plasma corticosterone to the log of plasma ACTH was calculated as an index of adrenal sensitivity to the action of ACTH (Engeland et al., 1981; Jasper and Engeland, 1997) for each time point.

#### 3.3.2 Experiment 2

##### Delayed spatial win shift task (DSWS)

A separate cohort of animals underwent an 11-week recovery period (males) or a 15-week recovery period (females) following a 3-week chronic variable stress (CVS) protocol. The delayed spatial win-shift (DSWS) task was adapted from (Wenk, 2004). After recovery, rats completed a standardized habituation sequence. On day 1, animals were switched to a reverse light cycle. Three days later, they were acclimated to the red-light testing room for 1 hour/day. On day 5, animals began food restriction to 70% of baseline intake, maintained throughout the experiment.

Rats were then habituated to the sucrose reward by receiving four sucrose pellets/day for three days. Subsequently, animals were allowed 10 minutes of free exploration in an 8-arm radial maze, followed the next day by another 10-minute session with sucrose pellets placed at the end of each arm. Formal DSWS testing began the next day and lasted 12 days, consisting of training, delay, and test phases. During training, rats were given up to 5 minutes to explore the maze, where four arms were blocked and four were baited. After visiting all baited arms, animals were returned to their home cage for a 5-minute delay period. In the subsequent test phase, the previously blocked arms were the only baited arms. Rats had up to 5 minutes to visit all four baited arms, using spatial cues in the experimental room. Baited arm assignments were randomized daily. The maze was cleaned with 20% ethanol between animals. Training and testing continued daily until animals reached criterion, defined as visiting all four baited arms in five or fewer choices on two consecutive days. Errors were categorized as within-phase (re-entries into previously visited test-phase arms) or across-phase (entries into arms baited during training).

#### 3.3.4 Experiment 3

##### Novel object recognition (NOR)

After 5 weeks of recovery from the 2-week CVS protocol, a new cohort of rats was tested in the novel object recognition (NOR) task (adapted from (Bevins and Besheer, 2006)). On day 1, animals were acclimated to the testing room for 2 hours and then habituated to the arena for 10 minutes. Following habituation, two identical objects were placed in the arena, and each rat was reintroduced facing away from the objects for a 10-minute sample phase (Familiarization 1). Animals were then returned to their home cages until the next day. On day 2, rats underwent a second familiarization session under the same conditions. After this session, animals were removed for 1 hour while the arena was cleaned with 70% ethanol. One familiar object was then replaced with a novel object. Rats were returned to the arena facing away from both objects and allowed to explore for 5 minutes. Placement of the novel object was counterbalanced across animals. Behavior was recorded using EthoVision XT (Noldus). Object exploration was defined as the rat’s nose entering a 2-cm radius around an object. Total exploration time was used to calculate the discrimination index: time investigating the novel object divided by the total time spent interacting with both objects. Rats that did not explore one or both sample objects during familiarization were excluded from analysis.

##### Morris water maze (MWM)

One week after NOR, the same group of rats was evaluated in the MWM to assess hippocampal-dependent spatial learning. The protocol was adapted from (Vorhees and Williams, 2006; O’Mahony et al., 2014). Animals were trained (acquisition days 1–4) with a hidden clear Plexiglas platform in a constant position located in one quadrant of the pool. Animals were subjected to four trials on each training day. For each trial the starting point varied randomly. Trials started with the rat facing the wall of the pool. The animal was released into the water at water-level. Time was started the moment the animal was released and measured until the rat located the submerged platform. On conditions where the animal was unable to locate the platform within the allocated time frame (95 seconds), it was then guided to the platform by the experimenter, where they had to wait for 20 s before being rescued. Cages were run randomly during the assay. During training, data was collected manually with a stopwatch by a blind operator looking to the video recording system. For the probe trial (day 5) the platform was removed from the pool and the animal was placed in a novel start position in the maze, facing the tank wall. The animal was then removed after 60 seconds. One animal was removed from analysis for considering as non-performer, after consecutively failing to find the platform, and when guided to it, failed to remain on the platform for the required time to reach criteria to finish the session. Videos from the probe session were tracked using EthovisionXT, Noldus.

#### 3.3.4 Experiment 4

##### 3-chamber social test

Preference for social novelty was assessed in same group 48 h later, as described previously (O’Tuathaigh et al., 2007; Yang et al., 2011). Briefly, rats were placed in a rectangular apparatus (80×40×40cm) divided into 3 equal-size chambers by transparent partitions each with a small opening (door) allowing easy access to all compartments. The test was composed of 3 sequential 10 minute trials; trial 1: habituation (the rat was allowed to explore the 3 chambers, confinement cylinders were empty), trial 2: sociability (an unfamiliar rat was placed into one of the cylinders in either the left or right chambers and exploration of the 3 chambers by the experimental rat was recorded for a further 10 minutes), trial 3: social novelty preference (a novel second rat was placed into the cylinder in the chamber opposite the (now familiar) rat from the previous stage. Exploration of the 3 chambers by the test rat was again recorded for 10 minutes. Stimulus rats used in this test were younger Sprague Dawley rats and they were housed in a different room than the experimental animals. Between each trial the experimental animal was removed, and each chamber was cleaned with 70% ethanol between. For each of the 3 stages, behaviors were recorded by a video camera mounted above the apparatus, and the EthovisionXT (Noldus) automated tracking system was used to analyze the time spent and the frequency of entry into each of the chambers. Social interaction index was calculated as percent of time spent on the social chamber/total end chambers time. The threshold criterion for discrimination in this phase was 60%. In the social memory phase, recognition index was calculated as percent of total time in the novel animal chamber/total time interacting with both animals. Since the mean value of all the groups was barely over 60% the discrimination criterion was reduced to 55%. Two animals (one of each sex) that had 0 time in any of the target chambers and remain immobile in a corner during the testing phases were as considered non-performers and removed from analysis.

##### Passive Avoidance

Lastly, 4 days after, these rats were evaluated in the passive avoidance test following protocol from (Martelle et al., 2021). The automated Active and Passive Avoidance System passive avoidant apparatus consisted of two, equal size chambers (24 cm (W) x 20cm (D) x 20cm (H)) separated by a guillotine door (GEMINI, San Diego Instrumentals, USA). One of the chambers was illuminated, while the other remained dark. On habituation day, the rat was placed into the illuminated chamber. After 30 s, the guillotine door was raised, and the rat was able to explore for 120 s. Latency to cross into the dark chamber and number of crosses between chambers were scored. A day after, the training phase consisted of a 30 s habituation to the illuminated chamber before the guillotine door was raised. This time, once the rat crossed to the dark chamber the guillotine door closed and the rat received a 0.25 mA shock during 1 s. The rat remained in the dark chamber for 10 s after the shock. On test day, 24 hours later, after the door open (30s from the onset) the trial continued for a max of 300 s in which the latency to cross to the dark chamber was scored. In case the rat crossed to the other side, no shock was delivered, and the rat remained in that chamber for 10 second before returning to their home cage. Finally, a retention test identical to the original test day was given 1 weeks later.

#### 3.3.5 Experiment 5

##### mRNA expression analysis

In order to determine possible signaling systems involved in the generation of the effects of chronic adolescent stress, we followed the expression trajectory of several mRNA targets of interest immediately after the cessation of the CVS protocol and 5 weeks after, the typical recovery time applied in most of our experiments. Some of these specific genes were chosen based on their well-known function on stress signaling and in other cases they were selected based on preliminary adult chronic stress results from our lab (unpublished data). A new group of animals subjected to adolescent CVS (as in experiments 3 and 4) and their respective controls were used. Rats were rapidly decapitated, and the brains were quickly removed and immediately flash-frozen in isopentane cooled in dry ice. IL and BLA were dissected from flash-frozen brains on cryostat (Microm HM550MP) at -16 °C based on morphological landmarks and coronal sections using an atlas (Paxinos and Watson, 2014). 500 μm sections were mounted on a chilled slide and bilateral tissue punches were obtained using a microdissection brain punch (Stoelting Company, Wood Dale, IL) with a diameter of 1 mm (IL) and 0.75 mm (BLA). Tissue was homogenized in lysis buffer and RNA was isolated using the RNA was extracted with Qiagen RNeasy Micro Kit. (REF74004) according to the manufacturer’s instructions. Prior to reverse transcriptase PCR, the RNA was treated with DNAse to remove any trace genomic contamination and cDNAs were synthesized employing SuperScript®IV First-Strand Synthesis System (Invitrogen, Life Technologies, cat 18091050). The purity and integrity of the RNA was assessed by spectrophotometry and agarose gel electrophoresis. Samples were diluted 1ul cDNA + 4ul H2O. Expression levels were evaluated by quantitative real-time PCR (qPCR) using TaqMan® Fast Advanced Master Mix (cat 4444557) and Custom Designed TaqMan Array Micro Fluidic Cards (Thermofisher). These gene represent pathways invoked or regulated during chronic stress, including glucocorticoid receptor (Herman et al., 2020), adrenergic (Kvetnansky et al., 2009), serotonin (Chaouloff et al., 1999), GABAergic signaling (Maguire, 2014), stress-related neuropeptide (CRH (Herman et al., 2020), PACAP (adcyap1)(Boucher et al., 2021), proenkephalin (Henry et al., 2017)) and neuroplasticity signaling pathways (Egr1 (Cullinan et al., 1995a; Gallo et al., 2018), BDNF (Girotti et al., 2024)).

The housekeeping genes used were 18s, b-Actin and Hprt1. Data were normalized against the geometric mean of their CT values. Results were analyzed comparing ΔCT values. Relative expression pattern was quantified using the ΔΔ method.

#### 3.3.6 Statistical analysis

Depending on the experiment, analysis of variance (ANOVA) (Stress x Sex), two tailed Student’s (control vs adol CVS), repeated measurements ANOVA (adol CVS x Time; adol CVS x Object; or Adol CVS Chamber) were used for statistical analysis with a level of significance of p≤0.05. In tests were preference or recognition indexes were calculated, we also performed one sample t tests against the threshold criteria for discrimination between the targets. In the cases where significant interactions were found on ANOVA tests, we performed the Šídák post-hoc analysis. In some cases, when only a main effect was found, we performed planned comparisons to evaluate individual differences between groups. In case of non-Gaussian distribution or heteroscedastic data, appropriate transformations were applied in ANOVA analyses. In the case of t tests, the Welch’s, t(W,) correction was used for non-homogeneous variance, Mann-Whitney U test for non-parametric data and Kolgomorov-Smirnov (KS) test for both heteroscedastic and non-parametric. F and t values with corresponding degrees of freedom expressed in each case and all graphical data is presented as mean values ± SEM except mRNA data that is presented as violin plots of fold change (FC) of adol CVS groups compared to the reference of mean FC value of control group (100%).

The main goal of this study was to analyze the effects of the combination of adolescent CVS in both males and females focusing on the specific effects within each sex. Considering that, and the great number of animals used for each experiment, it was logistically complex to carry out all the behavioral evaluations and to process all the tissue samples simultaneously to consider sex as a statistical variable. Therefore, male and female results were analyzed independently and adol CVS effects can only be interpreted within each sex. Data were analyzed and graphed using Prism 10 (GraphPad Software, La Jolla California USA).

AI editing for style and grammar was performed using Copilot and ChatGPT.

## 4. Results

### 4.1 Experiment 1: Cued fear conditioning and HPA axis reactivity assessment

To assess PFC and general corticolimbic functional integrity post conditioning, we examined the retention of extinction memory in a fear conditioning paradigm. Retention of extinction is heavily reliant on infralimbic function and amygdala connectivity (Giustino and Maren, 2015; Bloodgood et al., 2018). Animals underwent a 3-week chronic variable stress (CVS) protocol from postnatal day 40 ± 2 to postnatal day 61 ± 2. Restraint was excluded as a stressor to later assess the hypothalamic-pituitary-adrenal (HPA) axis response to a novel stressor. Sexes were analyzed separately in this experiment. After a 5-week post-CVS recovery period, they were tested in a cued fear conditioning paradigm. Two weeks post-conditioning, stress response reactivity was evaluated after exposure to a novel acute stressor (30-min restraint) (Timeline Fig. 1-A). Figures 1.B-C show the performance of the animals on the different phases of fear conditioning. Neither male nor female rats showed effect of adolescent CVS on their freezing levels during this test. The curves for each of the phases had a significant effect of Time (Male rats: Conditioning: F_(4, 72)_= 64.96, p≤0.0001; Extinction: F_(4, 72)_= 26.57, p≤0.0001; Recall: F_(2, 36)_= 11.21, p= 0.0002. Female rats: Conditioning: F_(4, 72)_= 41.19, p≤0.0001; Extinction: F_(4, 72)_= 42.58, p≤0.0001; Recall: F_(2, 36)_= 29.72, p≤0.0001). Two weeks later, when these animals were exposed to a novel acute stressor (restraint 30 min), female rats subjected to adol CVS had decreased levels of ACTH, adol CVS F_(1, 18)_ = 8.236, p= 0.01 as well as an adol CVS x Time interaction F_(4, 72)_ = 3.47, p= 0.01 (Time effect F_(4, 72)_ = 38.18, p≤0.0001). The post hoc analysis revealed that the group submitted to prior adol CVS had reduced ACTH secretion both at 15 and 30 minutes after the onset of restraint (p≤0.05 respectively)(Fig. 1-G). There was no main effect of adol CVS when analyzing corticosterone levels, but there was a significant interaction over time (Time F_(4, 72)_ = 106.5, p≤0.0001; adol CVS x Time F_(4, 72)_= 9.395, P≤0.0001). Female rats with previous adol CVS secreted less corticosterone 60 minutes after the onset of the novel stressor (Fig- 1-H). When analyzing the index of adrenal sensitivity (Jasper and Engeland, 1997), calculated as the concentration of corticosterone over the logarithm of ACTH concentration for each time point, we get an idea of how responsive the adrenal cortex was to effects of ACTH across the time we evaluated the HPA axis response to the acute stressor. We observed a shift in sensitivity between the peak and the recovery phase of the response in female rats. There was a main effect of Time F _(4, 72)_ = 81.17, p≤0.0001 and Adol CVS x Time interaction F _(4, 72)_ = 9.469, p≤0.0001 and the post hoc analysis showed a significant increase in the adrenal sensitivity at 30 min and a significant decrease at 60 min (p≤0.05 respectively) (Fig- 1-H). In the case of males, there was only an effect of time in all the variables analyzed (ACTH F_(4, 72)_ = 38.98, p≤0.0001; Cort F_(4, 72)_ = 111.4, p≤0.0001, adrenal sensitivity F _(4, 72)_ = 109.2, p<0.0001 (Fig. 1D-F).

**Fig. 1:**
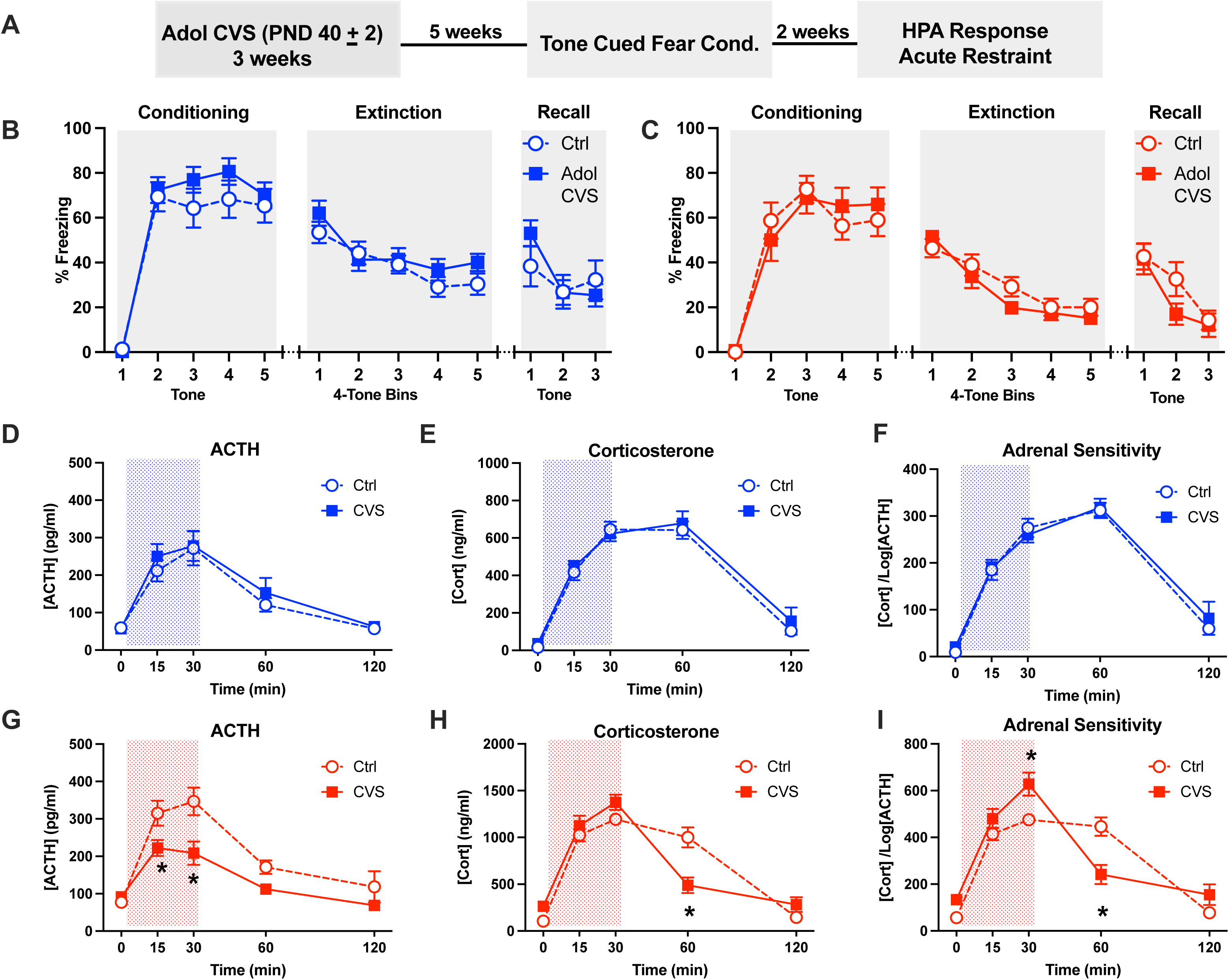
Experiment 1. Cued fear conditioning and HPA axis reactivity assessment: (A) <u>Experimental timeline.</u> (B-C) <u>Fear conditioning phases</u>. <u>HPA response to 30-min</u> <u>acute restraint:</u> ACTH (D, G), corticosterone (E, H), adrenal sensitivity (F, I). Male results in blue and female results in red. Shaded area represents time of restraining. Data presented as mean values ± SEM. N=10 each group. * p<0.05 against corresponding control.

### 4.2 Experiment 2: Delayed spatial win shift task (DSWS)

A second cohort from the same litters as Experiment 1 (timeline Fig. 2-A), was tested in the delayed spatial win-shift (DSWS) task, which relies on spatial working memory and executive control mediated by the medial prefrontal cortex and supported by hippocampal visuospatial processing (Floresco et al., 1997; Taylor et al., 2003; Wirt and Hyman, 2017) As in Experiment 1, significant effects were observed only in female rats. Females previously exposed to adolescent CVS showed an increased number of within-phase errors (t_(11)_ = 2.33, p = 0.038), with no differences in across-phase errors or total errors (Fig. 2-C). Across the 12-day protocol, within-phase errors showed main effects of adolescent CVS (F(1,18) = 5.44, p = 0.032) and time (F(11,198) = 3.99, p < 0.0001), but no CVS × time interaction. Planned comparisons indicated higher error rates on days 1 and 3 (p < 0.05) (Fig. 2E). For across-phase errors, there was a main effect of time (F(11,198) = 3.64, p < 0.0001) and a significant CVS × time interaction (F(11,198) = 2.01, p = 0.029). Šídák’s post hoc test showed that CVS females made more across-phase errors on day 1, but this deficit resolved thereafter (Fig. 2-G). Male rats showed no effects of adolescent CVS on any error measure (Fig. 2-B,D,F). Adolescent CVS did not affect acquisition time in either sex, as all groups reached criterion in a comparable number of days (Fig. 2-H,I).

**Fig. 2:**
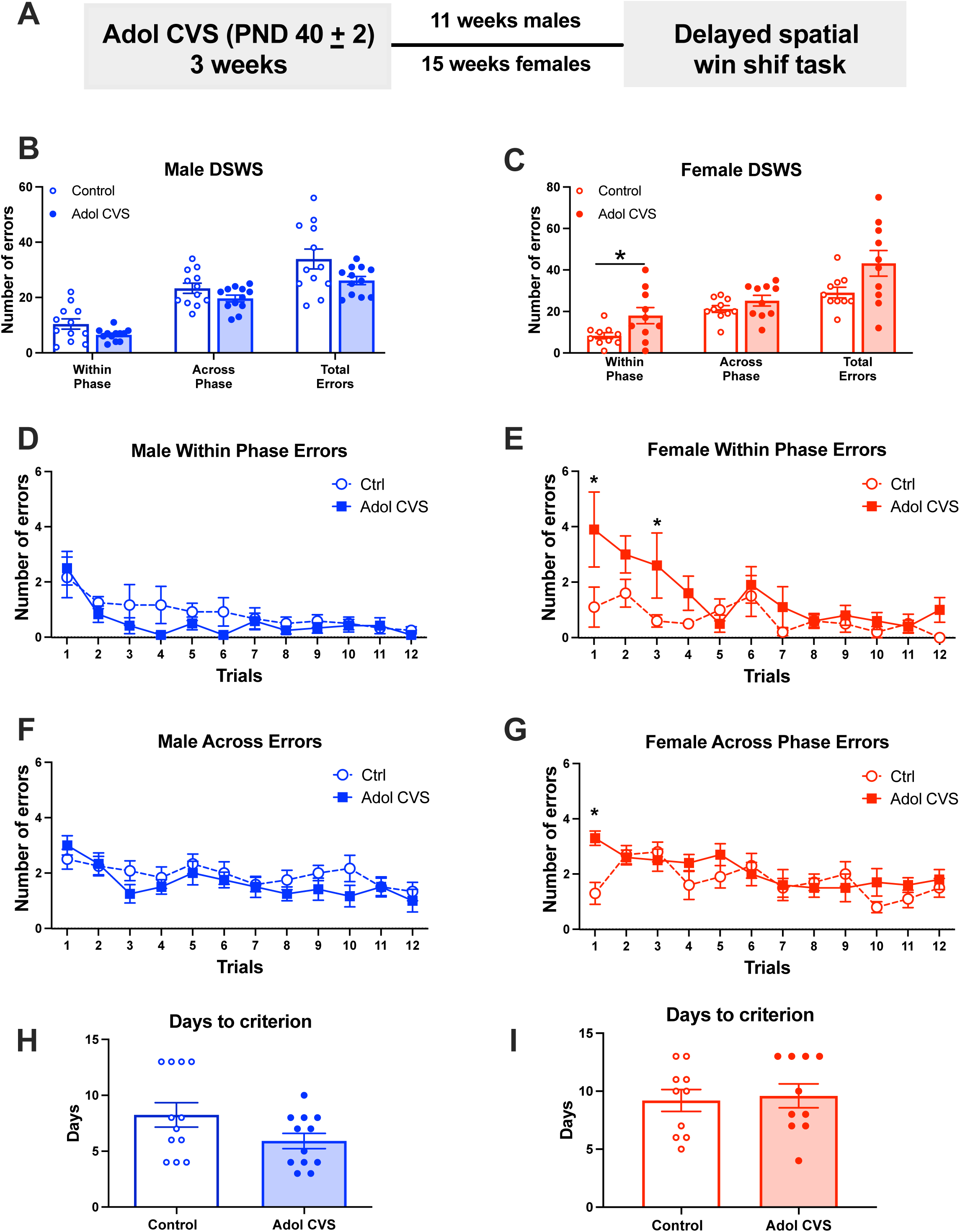
Experiment 2. Delayed spatial win shift task (DSWS): (A) <u>Experimental</u> <u>timeline</u>. (B-C) Average of each time of error. Within phase errors across training (D-E). Across phase errors across training (F-G). Days to reach criterion (H-I). Male results in blue and female results in red. Data presented as mean values ± SEM. N=10 each male group. * p<0.05 against corresponding control.

### 4.3 Experiment 3: Novel object recognition (NOR) test and visuo-spatial orientation in the Morris Water Maze (MWM)

To further assess cognitive functions involving recognition memory and visuo-spatial learning (Bevins and Besheer, 2006; Vorhees and Williams, 2006), a new group of rats was subjected to a 2-week protocol of CVS from PND 46 ±2 to PND 60 ±2 and then allowed to recover for 5 weeks before being tested for novel object recognition (NOR) followed by the Morris water maze a week later (Timeline Fig.3 A). There were no effects of adol CVS on the exploration time of the novel object. Control and adol CVS groups of both sexes s explored the novel object more than the familiar one (male group: Object F _(1, 14)_ = 26.37, p=0.0002, planned comparisons familiar vs novel: control p<0,001, adol CVS p<0,05; female group: Object F _(1, 15)_ = 64.86, p<0,0001, planned comparisons familiar vs novel, control p<0,0001, adol CVS p<0,001 (Fig. 3-B,D). Although there were no differences between treatments groups in either sex, a sample t test against the discrimination threshold of 60% indicated that the male adol CVS group did not differ (t_(7)_ =1.238, p=0.257) (, indicating less efficient discrimination than the male control group (t_(6)_ =4.004, p=0.01) (Fig. 3-C). In the case of females, both groups significantly discriminated the novel (control t_(8)_=8.187, p≤0.0001; adol CVS t_(7)_=13.28, p≤0.0001). (Fig. 3-E).

**Fig. 3:**
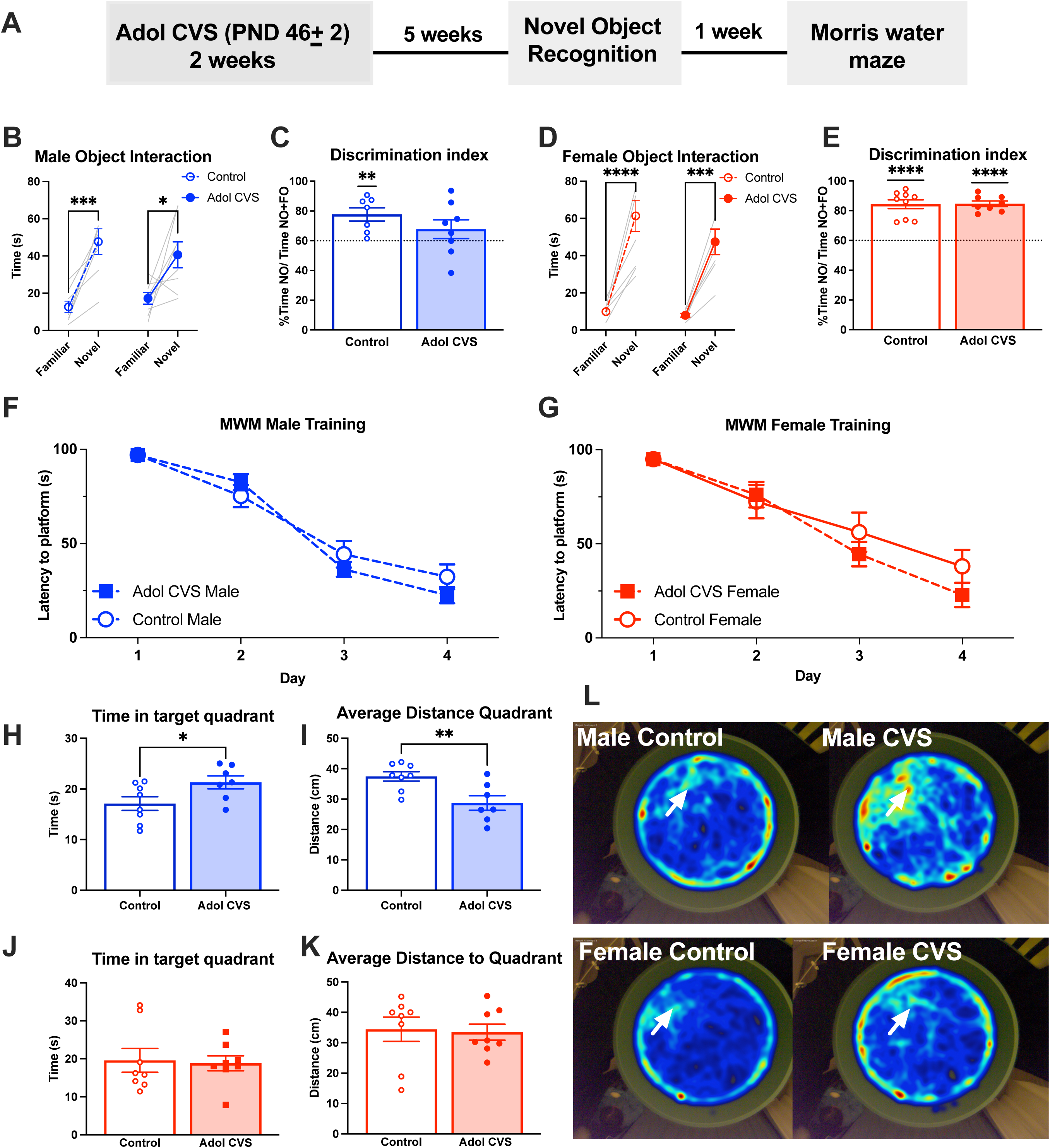
Experiment 3. Novel object recognition (NOR) test and visuo-spatial orientation in the Morris Water Maze (MWM): (A) <u>Experimental timeline</u>. <u>NOR:</u> (B-E) Total time of interaction with objects (* p<0.05, *** p<=0.001, **** p<=0.0001 vs respective control, two-way RM ANOVA) and discrimination index (<u>**</u> p<0.01, <u>****</u> p<=0.0001, vs criteria (dotted line), one sample t-test). <u>MWM:</u> (F-G) Training across sessions (average of 4 trials each day). (H-K) Probe test. Male results in blue and female results in red. Data presented as mean values ± SEM. (L) Merged heatmaps of all individuals in each group. White arrows indicate previous location of the platform.

A week after NOR testing, animals were evaluated in the Morris Water Maze. There were no effects of adolescent CVS on acquisition in either sex during the training phase (Fig. 4-A-B). In the probe test, however, males exposed to adolescent CVS showed enhanced memory for the platform location, demonstrated by increased time in the target quadrant (t(13) = 2.233, p = 0.0437) and a shorter average distance to the probe quadrant (t(13) = 3.148, p = 0.0077) (Fig. 4-C-D). No such effects were observed in females (Fig. 4-E-F). Adolescent CVS did not affect latency to reach the probe quadrant or the number of entries into the probe quadrant in either sex (not shown).

**Fig. 4:**
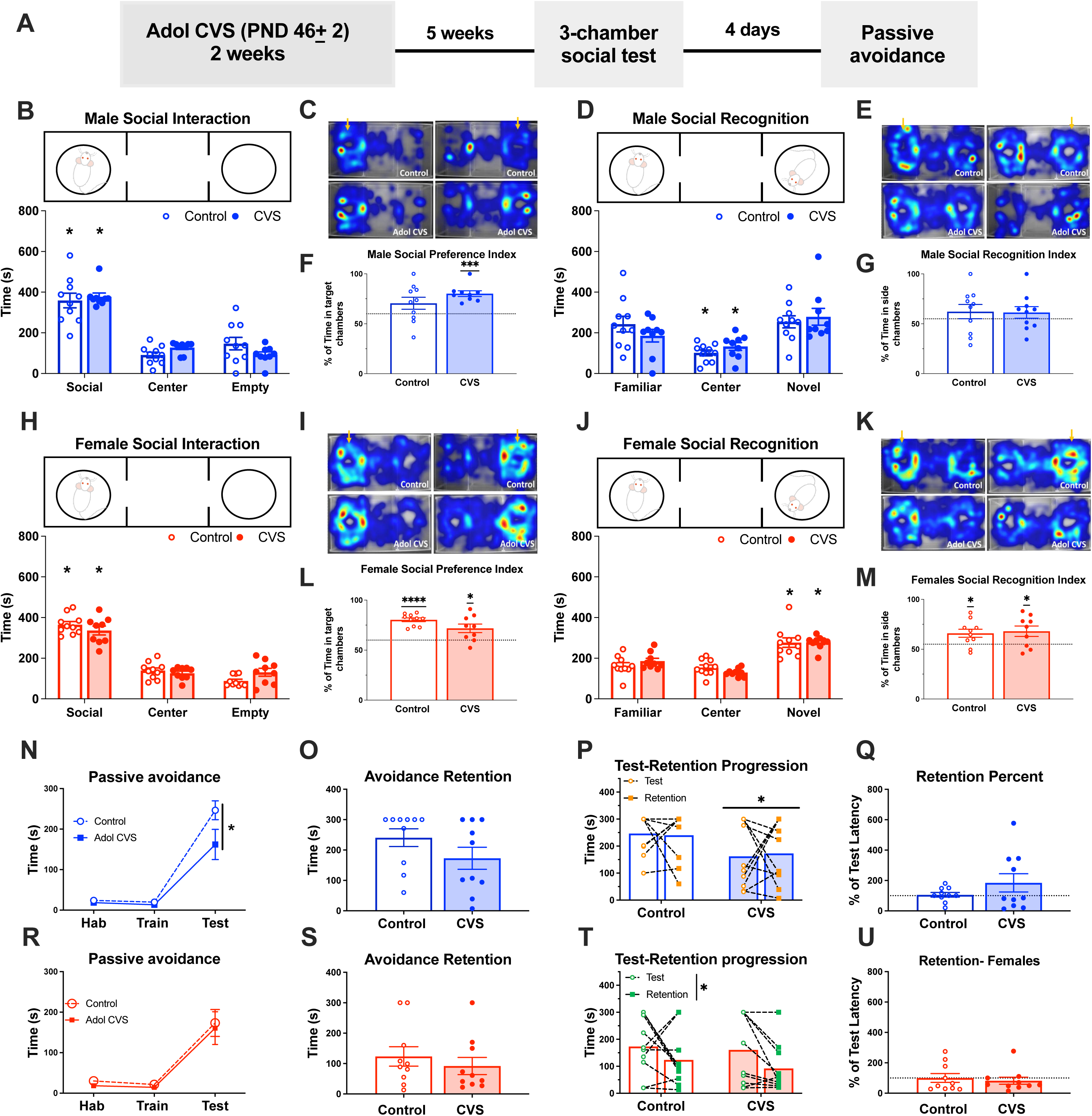
Experiment 4: 3-chamber social test and passive avoidance: (A) <u>Experimental timeline</u>. <u>Social preference:</u> (B, H) Time in social chamber vs other chambers (* p<0.05 vs all other chambers). (F, L) Social preference index (<u>*</u> p<0.05, <u>***</u> p<=0.00, <u>****</u> p<=0.0001, vs criteria (dotted line), one sample t-test). <u>Social memory.</u> Social recognition: (D, J) Time in chamber with novel animal vs other chambers (* p<0.05 vs all other chambers). (G, M) Social recognition index ((<u>*</u> p<0.05 vs criteria (dotted line), one sample t-test) Heat maps: Merged of all animals in each group, presented with novel social cue either left or right. Yellow arrows represent chamber where the novel social cue was placed in each case. <u>Passive avoidance:</u> (N, R) Phases of 3-day training test paradigm (* p<0.05 vs corresponding control). (O, S) Delayed retention 8 days later. (P, T) Individual trajectories between test and retention days (<u>*</u> p<0.05 vs control, |* p<0.05 between test phases). (Q, U) Percent retention/test day. Male results in blue and female results in red. Data presented as mean values ± SEM.

### 4.4 Experiment 4: Social preference, short-term social memory, and passive avoidance

A new cohort of animals was evaluated for social cognition aspects in the social 3-chamber test (social preference and social recognition) and passive avoidance to assess additional cognitive domains dependent on the medial prefrontal cortex, amygdala, and hippocampus (Torres-García et al., 2017; Yang and Wang, 2017; Tzakis and Holahan, 2019). After 2 weeks of adolescent CVS (PND 46 ± 2 to PND 60 ± 2) and 5 weeks of recovery (Timeline Fig. 4-A), animals were tested in the social preference and social recognition tasks, followed four days later by passive avoidance.

#### Social preference

In the preference phase, both male and female rats, regardless of treatment, spent significantly more time in the social chamber than in the other chambers (male: F(2,34) = 52.78, p < 0.0001; female: F(2,34) = 103.1, p < 0.0001; planned comparisons, p < 0.05) (Fig. 1- B,H). The social preference index did not differ between control and CVS groups in either sex. However, when compared to the discrimination threshold, only CVS males exceeded the threshold (t(8) = 7.12, p < 0.0001), whereas control males did not (t(9) = 1.76, p = 0.113), indicating a stronger preference in CVS males (Fig. 4- F, L).

#### Social recognition

When a novel animal was introduced, male rats from both groups showed no discrimination between the familiar and novel stimulus. Although there was a main effect of chamber (F(2,34) = 8.15, p < 0.001), post hoc analysis showed this reflected increased time in the center chamber only (p < 0.05) (Fig. 4-D). Discrimination index scores confirmed the absence of recognition in males (control: t(9) = 1.02, p = 0.33; CVS: t(9) = 1.105, p = 0.30) (Fig. 4-G).

In contrast, female rats, irrespective of CVS exposure, spent significantly more time with the novel animal (F(2,34) = 30.35, p < 0.001; planned comparisons, p < 0.05), and both groups exceeded the discrimination threshold (control: t(9) = 2.70, p = 0.024; CVS: t(8) = 2.5, p = 0.038) (Fig. 4-M).

#### Passive avoidance

In males, adolescent CVS significantly affected performance of passive avoidance (CVS: F(1,18) = 4.420, p < 0.05; phase: F(2,36) = 72.40, p < 0.0001; interaction: F(2,36) = 3.254, p < 0.05). CVS males had reduced latency to enter the dark chamber during the test phase, with no differences in other phases (Fig. 4-N). Females showed no CVS effects (phase: F(2,36) = 29.10, p < 0.0001) (Fig. 4-R). Eight days later, retention of avoidance memory was assessed (Fig. 4-O). Despite reduced latency in the test phase, CVS males did not differ from controls during retention (t(18) = 1.451, p = 0.16). A repeated-measures analysis across test and retention phases revealed a main effect of adolescent CVS (F(1,18) = 5.7, p = 0.028), reflecting overall reduced latency across phases, with no phase effect or interaction (Fig. 4-P). Inspection of individual trajectories indicated greater variability in the CVS group; percent change scores also showed higher dispersion, though not statistically significant (Fig. 4-Q). No effects were observed in females (Fig 4-R-U).

### 4.5 Experiment 5: mRNA expression analysis

We measured the expression of multiple target genes in the infralimbic (IL) and basolateral amygdala (BLA) immediately and 5 weeks after adol CVS. Differences were observed depending on brain region and time-point. Immediately after CVS we observed that in male rats, adol CVS evoked a reduction of reduction of CRH (t_(16)=_2.464, p= 0.0254) and BDNF (t_(18)_=2.516, p=0.0216) in the infralimbic cortex (Fig. 5-A, C), while 5 weeks after the expression of those genes was normalized and only Fkbp5 was upregulated (KS D_(n=10-9)_= 0.7778, p= 0.0065 (Fig. 5-B, C). In female rats, immediately after adol CVS we observed a reduction in the mRNA levels of two components of GABAergic signaling, the receptor subunit gamma-1 (Gabrg1 or GABAA γ1) t_(18)_=2.143, p= 0.0460, and the GABA transporter 3 (Slc6a11 or GAT-3) t_(17)_=2,369, p= 0.03. 0192, as well as a reduction in the opioid precursor proenkephalin (Penk) t(17)=2.195, p= 0,0423 (Fig. 5-D, E). After 5 weeks, the GAT-3 transporter was upregulated t_(18)_=2.342, p= 0.0309, as well as the Beta1 adrenoreceptor (Adrb1) t(18)=2.535, p= 0.0207, and the early growth response protein 1 (Egr1 or ZNF268) t_(18)_=2.336, p= 0.0313 (Fig. 5- F, E).

**Fig. 5.**
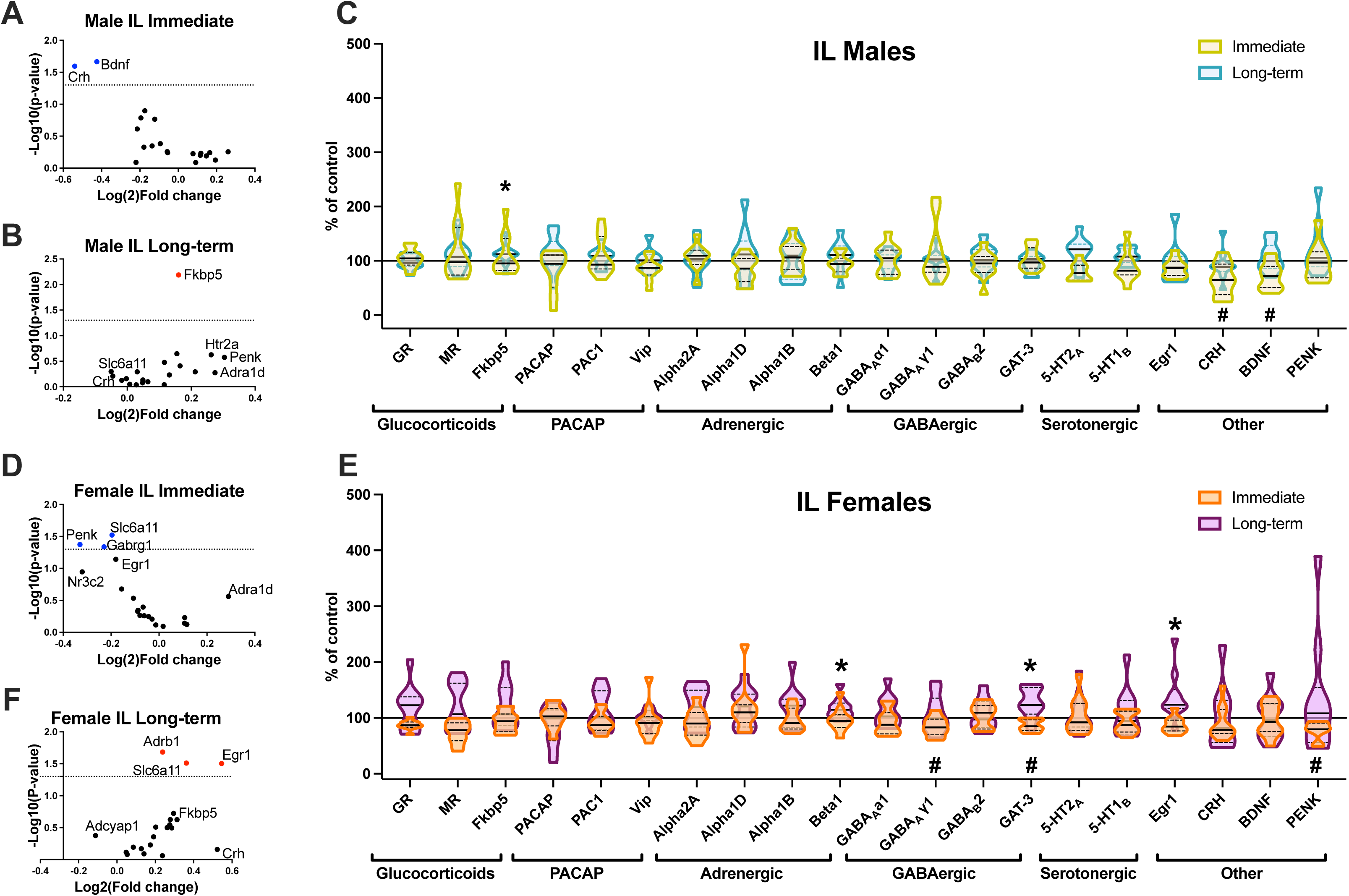
Experiment 5: mRNA expression analysis IL: Relative expression pattern of mRNA of target genes quantified immediately after adol CVS (2-week CVS protocol starting PND 46±2) of after a 5-week recovery period (Long-term). (A, B, D, F) Results for each sex and timepoint are displayed as volcano plots (dotted line at nominal p=0.05). Red dots=upregulated targets, blue dots=downregulated targets). (C, E) Overlapped relative expression for each time point expressed a percent from respective control group (mean of control=100%). # p<0.05 immediate effect. * p<0.05 long-term effect.

In the (BLA) of male rats, adol CVS caused increased beta1 adrenergic receptor (Adrb1) t_(17)_=2.251, p= 0.0379 and a reduction in serotonin receptor 1B (Htr1b or 5-HT1_B_) t_(W, 12.84)_=2.634, p= 0.0208 (Fig 6-A, C). Immediate changes were reversed 5 weeks later, but a late-emerging downregulation of the Adenylate Cyclase Activating Polypeptide 1 (Adcyap1,PACAP) t_(16)_=2.227, p= 0.0407, the alpha 1D adrenergic receptor (Adra1d, Alpha1D) t_(W, 12.84)_=2.676, p=0.0192 and proenkephalin (Penk) t_(14)_=3.013, p= 0.0093 were observed (Fig. 5-B, C).

**Fig. 6.**
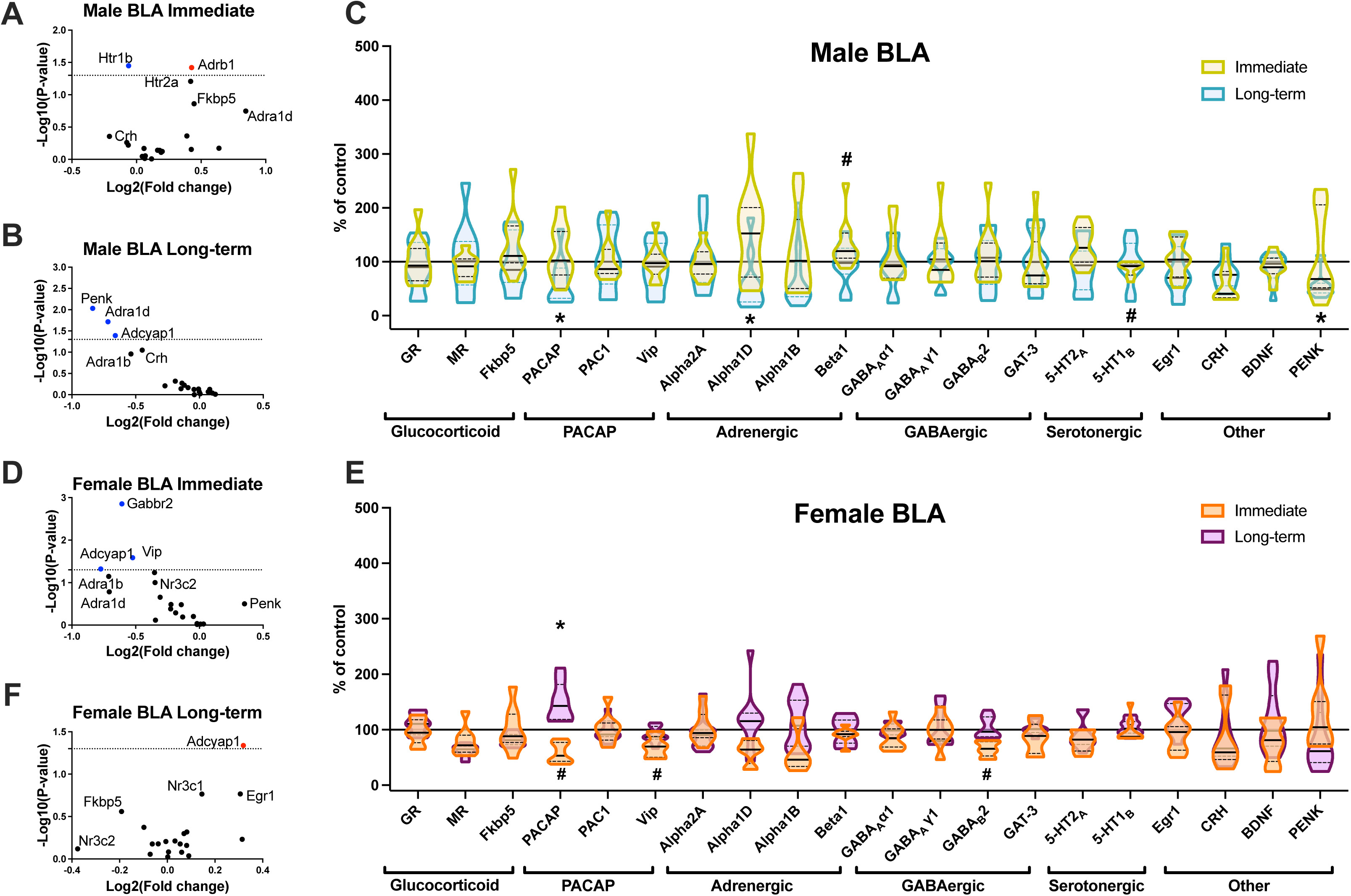
Experiment 5: mRNA expression analysis BLA: Relative expression pattern of mRNA of target genes quantified immediately after adol CVS (2-week CVS protocol starting PND 46±2) of after a 5-week recovery period (Long-term). (A, B, D, F) Results for each sex and timepoint are displayed as volcano plots (dotted line at nominal p=0.05). Red dots=upregulated targets, blue dots=downregulated targets). (C, E) Overlapped relative expression for each time point expressed a percent from respective control group (mean of control=100%). # p<0.05 immediate effect. * p<0.05 long-term effect.

In females, there was reduction in expression of PACAP (t_(17)_=2.133, p= 0.0478), VIP (t_(17)_=2.441, p= 0.0259) and GABAB2 t_(17)_=3.801, p=0.0014 mRNAs immediately after CVS (Fig. 6-D, E). While the two other genes VIP and GABAB2 returned to pre-stress levels, PACAP expression was now upregulated t_(14)_=3,472, p= 0.0037 (Fig. 6-F, E). Together, the post-stress changes in the expression of Fkbp5, PACAP, proenkephalin, and α1D- adrenergic receptor in male rats, and in PACAP, β1-adrenergic, GAT-3, Egr1 in female rats, reflect coordinated, long-lasting modulation of neuromodulatory systems that regulate excitability and stress responsiveness within IL–BLA circuits.

## 5. Discussion

Across behavioral, hormonal, and molecular endpoints, our data indicate that adolescent chronic variable stress (CVS) produces long-lasting, sex-specific adaptations that are not uniformly deleterious. Rather than inducing global cognitive or emotional dysfunction, adolescent CVS leads to selective alterations in stress regulation, prefrontal–amygdala-dependent behaviors, and neuromodulatory gene expression, with distinct profiles in males and females. These findings support the idea that developmental CVS reshapes, rather than uniformly disrupts, corticolimbic maturation, leading to distinct behavioral and molecular effects in males and females. Adolescence is a critical developmental window during which environmental conditions can exert lasting effects on brain circuitry. Prior work suggests that exposure to adversity during this period can promote divergent developmental trajectories, resulting in differential behavioral and neurochemical outcomes (Green and McCormick, 2013; McCormick and Green, 2013; Sheth et al., 2017). In the present study, we report a series of experiments examining the enduring consequences of adolescent CVS in both male and female rats.

Chronic or repeated stress during adolescence has been shown to exert long-term effects on multiple neural systems, including alterations in hippocampal morphology (Isgor et al., 2004), modulation of prefrontal and amygdala stress responsivity (Cotella et al., 2020), and lasting changes in hypothalamic–pituitary–adrenal (HPA) axis function in adulthood (Pohl et al., 2007; Bourke and Neigh, 2011; Wulsin et al., 2016; Cotella et al., 2019). In many cases, these adaptations may reflect stress resilience rather than vulnerability. For example, our prior work suggests that adolescent CVS in males promotes resilience to subsequent traumatic stress later in life (Cotella et al., 2023), reducing extinction impairments in a cued fear conditioning paradigm. In contrast to adult CVS, which produces lasting sensitization of the HPA axis response to restraint, adolescent CVS does not induce such sensitization (Cotella et al., 2019, 2020), further supporting the concept of neuroendocrine resilience.

Consistent with this framework, lasting changes in HPA axis function were observed in females exposed to adolescent CVS, characterized by reduced ACTH and corticosterone responses to novelty stress and altered adrenal sensitivity. These findings replicate prior work from our laboratory, indicating that this represents a reliable outcome of this exposure paradigm. Reduced stress reactivity occurred in the context of increased immobility in the forced swim test (Wulsin et al., 2016), suggesting engagement of alternative or passive coping strategies. In contrast, males exhibited no lasting changes in acute stress hormone responses, consistent with prior evidence of HPA axis resilience following adolescent stress.

Behaviorally, females exhibited mild cognitive alterations following adolescent CVS in the delayed spatial win-shift task, mainly due to an increase in within phase errors. This pattern is consistent with reduced efficiency of spatial working memory, a behavior linked to medial prefrontal cortex function (Taylor et al., 2003; Gisquet-Verrier and Delatour, 2006; Bizon et al., 2012). In contrast, females did not show deficits in passive avoidance or fear extinction, suggesting that adolescent CVS did not produce a broad impairment across all prefrontal-dependent behaviors (Gilmartin et al., 2014; Pellman and Kim, 2016; Marek et al., 2019; López-Moraga et al., 2022; Olmedo-Córdoba et al., 2023). Behaviors more hippocampal-dependent, like spatial memory in the Morris water maze (Terry, 2009; Othman et al., 2022), and social recognition memory (Meira et al., 2018; Tzakis and Holahan, 2019), as well as amygdala-biased behaviors (e.g., acquisition of conditioned fear (Pellman and Kim, 2016)) were unaffected, suggesting that the impact of adolescent stress in females may be directed toward specific prefrontal circuitry. Notably, females showed long-term alterations in infralimbic gene expression of genes reported to be involved in stress adaptations, that are related to β-adrenergic signaling (Ramos and Arnsten, 2007; Kvetnansky et al., 2009), neuroplasticity (Egr1) (Cullinan et al., 1995b; Duclot and Kabbaj, 2017; Gallo et al., 2018), and GABA clearance (Maguire, 2014; Scimemi, 2014), whereas no comparable changes were detected in the basolateral amygdala (BLA), where only PACAP, a peptide tightly involved in modulation of behavioral stress responses (Hammack and May, 2015) was increased following a transient downregulation after adol CVS. These findings are consistent with sustained stress-induced reorganization mainly of prefrontal information processing. Importantly, adolescent stress did not alter fear acquisition or extinction, arguing against a generalized disruption of prefrontal function or corticolimbic circuitry.

In contrast, males exhibited distinct behavioral phenotypes following adolescent CVS, including enhanced retention in the Morris water maze probe trial, impaired novel object recognition, and impaired retention in the passive avoidance task. Fear conditioning remained unaffected, indicating intact Pavlovian learning. This suggests that the passive avoidance phenotype observed may reflect altered inhibitory avoidance performance or weaker response inhibition, rather than a general deficit in aversive learning, considering passive avoidance relies on context conditioning learning (Roesler et al., 2003; McAllister-Williams et al., 2010). Enhanced performance in the probe test of the MWM is consistent with previous reports showing certain forms of juvenile or adolescent stress improved aspects of spatial memory tested this test (Frisone et al., 2002; Pansarim et al., 2023). Together, these outcomes are consistent with adolescent stress-related reorganization of prefrontal and hippocampal circuitry involved in spatial memory and avoidance.

The passive avoidance data warrant additional consideration. Analysis of behavior across the test and one-week retention sessions revealed substantial variability in male performance, suggesting the presence of distinct inhibitory response profiles. This variability likely reflects different behavioral endophenotypes that interact with prior stress exposure to bias animals toward alternative strategies (Wood et al., 2010; Calvo et al., 2011; Zurawek et al., 2019). In contrast to females, which showed a relatively uniform reduction in latency consistent with extinction of the inhibitory response, males did not exhibit a consistent trajectory. Some males appeared to extinguish the avoidance response, whereas others displayed increased latency at retention despite low latency during the initial test. This pattern indicates that, although impulsive responding was evident during testing, learning of the aversive contingency likely occurred. Larger, higher-powered studies will be required to better resolve these behavioral strategies and determine how developmental stress influences their prevalence.

Adolescent stress in males was associated with long-term increase of Fkbp5 in IL. This is a co-chaperone of the glucocorticoid receptor that can regulate intracellular GR sensitivity in response to stressors (Binder, 2009; Zannas et al., 2016; Malekpour et al., 2023), which may reflect a lasting recalibration of prefrontal GR signaling, potentially contributing to downstream regulation of PFC-dependent behavioral strategies as were observed in our experiments.

In the BLA of males, we observed a long-term baseline downregulation of PACAP, α1D-adrenergic receptors, and proenkephalin in the BLA. The former two, associated with excitatory processes (Ramos and Arnsten, 2007; Cho et al., 2012; Perez, 2020; Boucher et al., 2021), may be potentially contributing to reduced BLA drive, whereas changes in enkephalin expression may also favor altered inhibitory regulation within the BLA that may be associated with the behavioral adaptations observed. Penk reduction may be an interesting understudied molecular candidate to understand effects of the adol CVS model. Reduced enkephalin in the BLA has been associated with stress vulnerable phenotypes related to anxiety behavior (Bérubé et al., 2014). Together with our previous finding that males exposed to adol CVS showed increased anxiety-like behavior in the open field but reduced Fos recruitment in BLA, as well as other amygdala divisions, acutely after forced swim (Cotella et al., 2020), the present molecular data suggest that developmental stress may induce a long-term reorganization of amygdala responsivity rather than a simple shift in global amygdala drive. This interpretation is compatible with evidence that amygdala-mediated behaviors depend on heterogeneous microcircuits and projection-defined populations, whose activity can exert distinct, even opposing, effects on emotional behavior (Felix-Ortiz et al., 2015; Sharp, 2017; Daviu et al., 2019) In this context, long-term downregulation of PACAP, α1D adrenergic receptor, and proenkephalin in the BLA may reflect altered excitatory and opioid modulation of local BLA processing, potentially indicating stress-related mechanisms underlying altered BLA dependent behavioral effects. .

Sex-specific adol CVS effects were also evident in social behavior. Males showed little propensity to approach a novel conspecific in the three-chamber test, whereas stress-exposed males exhibited enhanced social preference to explore a novel conspecific. This effect may also be related to the dampened BLA output (Felix-Ortiz and Tye, 2014; Wellman et al., 2016) consistent with observed molecular adaptations. Even more, although not completely replicating our results, enkephalin (Bérubé et al., 2014) and PACAP (Schmidt et al., 2021) within the BLA have been reported to be involved in the processing of social behaviors.

We assessed stress-related gene expression in the infralimbic cortex (IL) and BLA immediately following cessation of a two-week adolescent CVS exposure and again five weeks later. In no case did the initial transcriptional changes persist into adulthood, and there was no overlap in differentially expressed genes across sexes. In females, immediate post-CVS reductions were observed in genes related to neuronal inhibition within the IL, including GAT-3, the GABA-A receptor γ1 subunit, and proenkephalin, suggesting reduced net inhibitory tone. In the female BLA, immediate reductions in PACAP, VIP, and GABA-B receptor subunits were observed, indicating a transient shift in excitatory–inhibitory balance. No differentially expressed genes from that list were detectable five weeks later, suggesting that these transcriptional changes were short-lived. However, the lack of persistent transcriptional differences on these genes at the later time point does not mean that these early changes were without functional significance. Even transient shifts in gene expression during adolescence may have lasting consequences on cellular development, synaptic organization, and circuit function.

In males, immediate post-CVS decreases were observed in genes associated with neuronal excitation and plasticity (e.g., 5-HT1B, CRH, BDNF), particularly within the IL, suggesting reduced excitatory processing. A smaller but significant reduction in 5-HT1B expression was observed in the BLA, indicating a more modest immediate impact in this region. Overall, these findings demonstrate that adolescent CVS differentially modulates gene expression across sexes and brain regions, with no evidence for persistent transcriptional changes into adulthood from that set of genes. Longer-term molecular adaptations were more apparent in males within the BLA and in females within the IL, suggesting region- and sex-specific reorganization of excitatory–inhibitory balance and neuroplasticity.

The gene set examined in this study was selected based on prior evidence of differential regulation in the IL and/or BLA following chronic stress, using targeted candidate-gene approaches. It is important to note that our findings do not fully align with transcriptome-wide analyses, which frequently identify substantial stress-related changes in non-neuronal cell populations, including oligodendrocytes, astrocytes, and microglia. Thus, the conclusions that can be drawn from the present study are necessarily limited. Future studies employing time-course designs combined with bulk or single-nucleus RNA sequencing will be required to fully resolve these molecular adaptations.

Our experimental design included both two- and three-week CVS regimens spanning late adolescence (postnatal days 40–61 or 46–60). Both regimens produced significant behavioral and endocrine phenotypes. Importantly, assessment of HPA axis responses following the three-week regimen replicated prior findings using a two-week stress exposure (Wulsin et al., 2016), confirming the sensitivity of this developmental window (Jankord et al., 2011). The persistence of adolescent stress effects is supported by behavioral phenotypes observed both five and eleven weeks post-CVS, although additional plasticity may occur during the intervening period.

In summary, adolescent chronic variable stress induces long-lasting, sex- and region-specific alterations in behavior, stress hormone regulation, and gene expression within prefrontal and amygdala circuits. Importantly, adolescent stress did not result in generalized cognitive deficits in adulthood. Spatial learning and recognition memory were largely preserved, and in some cases, such as Morris water maze performance, were enhanced, suggesting that certain forms of early stress can improve specific cognitive functions rather than being uniformly detrimental. In females, adolescent stress preferentially altered working memory and dampened hormonal responses to novelty stress, accompanied by transient shifts in genes related to GABAergic and adrenergic signaling. In males, adolescent stress produced variable inhibitory control in passive avoidance and altered expression of genes associated with stress signaling and plasticity (e.g., CRH, BDNF, FKBP5, PACAP). Collectively, these findings support the view that adolescent stress functions as a developmental pressure that shapes distinct patterns of resilience and vulnerability in males and females, rather than as an unequivocally pathogenic event.

## Acknowledgements

This project was funded by the National Institutes of Health (Grant Nos. R01MH101729, R01 MH049698, and R01 MH119814 [to JPH]; Grant No. T32 DK059803 [to EMC and SEM]; U.S. Department of Veterans Affairs (Grant No. I01BX003858 [to JPH]), and a NARSAD Young Investigator Award from the Brain and Behavior Research Foundation (to RDM).

Special thanks to Dr. Jessie Scheimann for assistance with the delayed spatial win shift test. We thank other members of Dr. Herman’s laboratory for their assistance in data collection and general discussion of the results.

The authors report no biomedical financial interests or potential conflicts of interest.

A previous version of this article was published as a preprint on bioRxiv.

